# “Heat waves” experienced during larval life have species-specific consequences on life-history traits and sexual development in anuran amphibians

**DOI:** 10.1101/2021.12.17.473144

**Authors:** János Ujszegi, Réka Bertalan, Nikolett Ujhegyi, Viktória Verebélyi, Edina Nemesházi, Zsanett Mikó, Andrea Kásler, Dávid Herczeg, Márk Szederkényi, Nóra Vili, Zoltán Gál, Orsolya I. Hoffmann, Veronika Bókony, Attila Hettyey

**Affiliations:** Lendület Evolutionary Ecology Research Group, Plant Protection Institute, Centre for Agricultural Research, Eötvös Loránd Research Network, Budapest, Hungary; Department of Systematic Zoology and Ecology, Eötvös Loránd University, Budapest, Hungary; Department of Ecology, Institute for Biology, University of Veterinary Medicine, Budapest, Hungary; Konrad Lorenz Institute of Ethology, Department of Interdisciplinary Life Sciences, University of Veterinary Medicine, Vienna, Austria; Doctoral School of Biology, Institute of Biology, Eötvös Loránd University, Budapest, Hungary; Animal Biotechnology Department, Institute of Genetics and Biotechnology, Hungarian University of Agriculture and Life Science, Gödöllő, Hungary

**Keywords:** *Batrachochytrium dendrobatidis*, Bufonidae, Chytridiomycosis, Heat stress, Ranidae, Thermal tolerance

## Abstract

Extreme temperatures during heat waves can induce mass-mortality events, but can also exert sublethal negative effects by compromising life-history traits and derailing sexual development. Ectothermic animals may, however, also benefit from increased temperatures via enhanced physiological performance and the suppression of cold-adapted pathogens. Therefore, it is crucial to address how the intensity and timing of naturally occurring or human-induced heat waves affect life-history traits and sexual development in amphibians, to predict future effects of climate change and to minimise risks arising from the application of elevated temperature in disease mitigation. We raised agile frog (*Rana dalmatina*) and common toad (*Bufo bufo*) tadpoles at 19 °C and exposed them to a simulated heat wave of 28 or 30 °C for six days during one of three ontogenetic periods (early, mid or late larval development). In agile frogs, exposure to 30 °C during early larval development increased mortality. Regardless of timing, all heat-treatments delayed metamorphosis, and exposure to 30 °C decreased body mass at metamorphosis. Furthermore, exposure to 30 °C during any period and to 28 °C late in development caused female-to-male sex reversal, skewing sex ratios strongly towards males. In common toads, high temperature only slightly decreased survival and did not influence phenotypic sex ratio, while it reduced metamorph mass and length of larval development. Juvenile body mass measured two months after metamorphosis was not adversely affected by temperature treatments in either species. Our results indicate that heat waves may have devastating effects on amphibian populations, and the severity of these negative consequences, and sensitivity can vary greatly between species and with the timing and intensity of heat. Finally, thermal treatments against cold-adapted pathogens have to be executed with caution, taking into account the thermo-sensitivity of the species and the life stage of animals to be treated.

## Introduction

Earth’s wildlife and ecosystem face the sixth mass extinction event today due to anthropogenic environmental alterations, including extreme climatic conditions (Ceballos et al. 2015). Due to global climate change, heat waves occur with increasing frequency, intensity and duration (Gardner et al. 2016). Extreme temperatures during heat waves expose species to intensified physiological stress (Williams et al. 2016) and can even induce mass-mortality events (Welbergen et al. 2008, McKechnie and Wolf 2019). Warming climate with frequently reappearing heat waves can alter species distributions (Krockenberger et al. 2012, Stillman 2019), trigger shifts in the timing of the breeding season and directly affect breeding success in a taxonomically diverse range of species (Blaustein et al. 2001, Oswald et al. 2008, Truebano et al. 2018, Stillman 2019). These factors can generate profound changes in community structure and ecosystem functioning via the formation of interactions between species with previously non-overlapping spatial or temporal distributions (Williams et al. 2016) and the alteration of predator-prey and host-pathogen systems (Blaustein et al. 2010, Cohen et al. 2019, Stillman 2019, Carreira et al. 2020). Fluctuations in temperature affect ectotherms in particular because they lack the metabolic, physiological and anatomical mechanisms that would allow them to maintain constant body temperature, and, therefore, ectothermsare able to maintain high physiological performance only within a narrower environmental temperature range than are endotherms (Clarke and Pörtner 2010).

Amphibians are among the most threatened vertebrate groups, because 41 % of the species are endangered (IUCN 2021), and almost 50 % show population declines worldwide, mainly due to anthropogenic environmental change (Stuart et al. 2004, Wake and Vredenburg 2008, Hof et al. 2011, Monastersky 2014, Campbell Grant et al. 2016). The growing incidence of meteorological extremes and rising temperatures resulting from global climate change and anthropogenic heat pollution (i.e. urban heat islands; Arnfield 2003, Brans et al. 2018) are major threats to amphibians. Their complex life cycle, usually including an aquatic stage, the unshelled eggs and a highly permeable integument make amphibians excessively sensitive to water availability. Also, though amphibian larvae generally exhibit a relatively high thermal tolerance (Ultsch et al. 1999, Sunday et al. 2011, but also see Harkey and Semlitsch 1988, Wallace and Wallace 2000, Bellakhal et al. 2014, Goldstein et al. 2017) temperatures as low as 30 °C experienced during the larval period can be detrimental to them. Heat can result in delayed metamorphosis (Goldstein et al. 2017), reduced body mass (Harkey and Semlitsch 1988, Phuge 2017, Lambert et al. 2018), disabled locomotor activity (Goldstein et al. 2017), sex reversal (Dournon et al. 1984, Wallace and Wallace 2000, Mikó et al. 2021) and biased sex ratios (Phuge 2017, Lambert et al. 2018, Ruiz-Garciá et al. 2021). Exposure of adult frogs to 30 °C or higher can increase stress hormone levels (Juráni et al. 1973, Narayan and Hero 2014) and enhance the processes that contribute to accelerated ageing (Burraco et al. 2020).

Emerging infectious diseases represent another serious threat to amphibians (Harvell et al. 2002, Pounds et al. 2006). Due to repeated introductions arising from human activities (Lips 2016, O’Hanlon et al. 2018), chytridiomycosis caused by the chytrid fungi *Batrachochytrium dendrobatidis* (*Bd*) and *Batrachochytrium salamandrivorans* (*Bsal*) (Van Rooij et al. 2015) has already led to the decline or extinction of several hundred species and continues to cause mass mortality events on five continents (Scheele et al. 2019). Since *Bsal* was only discovered eight years ago (Martel et al. 2013) and its known geographic distribution is much smaller (Spitzen-van der Sluijs et al. 2016), we focus here on the much better known and more widespread *Bd*. The fungus infects keratinous epidermal layers of the skin with waterborne motile zoospores (Berger et al. 1998), impairs its osmoregulatory function, which leads to shifts in electrolyte balance that can ultimately result in cardiac asystolic death in metamorphosed anurans (Voyles et al. 2009). Tadpoles exhibit keratinous elements only in their mouthparts; therefore they are less susceptible to *Bd* infection than subsequent life stages (Marantelli et al. 2004, Blaustein et al. 2005). Nonetheless, it is often the early ontogeny (larval and metamorphic stages) when individuals become infected, due to their aquatic lifestyle (Kilpatrick et al. 2010). The thermal optimum of this cold-adapted fungus is between 18-24 °C, and its growth ceases above 27-28 °C (Cohen et al. 2017, Voyles et al. 2017), while the vast majority of amphibian species can survive temperatures above 30 °C (Ultsch et al. 1999, Sunday et al. 2011). Consequently, when and wherever microclimatic conditions allow amphibians to sufficiently raise their body temperature via thermoregulation, *Bd* infection prevalence and intensity are low (Richards-Zawacki 2010, Forrest and Schlaepfer 2011, Becker et al. 2012), and mass mortalities typically only occur in constantly cool environments (Berger et al. 2004, Woodhams and Alford 2005). Accordingly, thermal treatment of amphibians with 28 °C and higher for a few days can be effectively applied for *Bd*-disinfection of larval, juvenile or adult amphibians in captive populations (Woodhams et al. 2003, Retallick and Miera 2007, Chatfield and Richards-Zawacki 2011, Geiger et al. 2011, McMahon et al. 2014) and may also prove effective for fighting *Bd in situ* (Hettyey et al. 2019).

Based on the above information, heat waves may exert several opposing effects on developing amphibians, which may be beneficial for combating *Bd* but harmful for other fitness-related traits. Heat waves usually last for only a couple of days, and just a few days of heat treatment can be sufficient for the suppression or even the complete clearance of cold-adapted pathogens, such as *Bd* (Woodhams et al. 2003, Retallick and Miera 2007, McMahon et al. 2014). However, we still know little about the developmental costs of brief periods of high temperatures for larval amphibians because most experiments that investigated the effects of heat on larval fitness exposed animals to heat chronically for several weeks, and the effects of shorter heat pulses are rarely tested (Mikó et al. 2021). Because the effects of high temperatures are likely to depend on the intensity, timing and duration of exposure, and may differ between species, studies focusing on these sources of variation are necessary to assess potential malign impacts of heat waves on amphibians and uncover hidden risks arising from thermal treatment of diseased animals.

In this study, we experimentally investigated the developmental effects of six-day long exposures to 28 and 30 °C during early, mid, and late larval development of two amphibian species. We assessed the effects of these experimental heat waves on the survival, growth, somatic and sexual development of agile frogs (*Rana dalmatina*; Bonaparte, 1840) and common toads (*Bufo bufo*; Linnaeus, 1758). These two species are common in Europe, they inhabit various types of water bodies, have different thermal optima (Morand et al. 1997) and represent two globally widespread families (Bufonidae and Ranidae). The temperatures we applied occur during heat waves in aquatic habitats of amphibian larvae in the temperate climate zone (Lambert et al. 2018, Lindauer et al. 2020) and are also recommended for thermal treatment of diseased amphibians (Chatfield and Richards-Zawacki 2011, McMahon et al. 2014, Cohen et al. 2017, Hettyey et al. 2019). Thus, our aim was twofold: to reveal developmental effects of heat waves that may occur in natural habitats, and to assess possible negative consequences of thermal treatment applied against cold-adapted pathogens.

## Methods

### Experimental procedures

In March 2019, we collected 50 eggs from each of 12 freshly laid egg clutches of the agile frog from three ponds in the Pilis-Visegrádi Mountains, Hungary (Katlan: 47.71110° N, 19.04570° E, Ilona-tó: 47.71326° N, 19.04050° E, and Apátkúti pisztrángos: 47.76656° N, 18.98121° E; four clutches from each population). We transported the eggs to the Experimental Station of the Plant Protection Institute in Julianna-major, Budapest, and placed each clutch (family hereafter) into a plastic container (24 × 16 ×13 cm) filled with 1.3 L continuously aerated reconstituted soft water (RSW; APHA et al. 1992, USEPA 2002). In the laboratory, we maintained 16.3 ± 0.3 °C (mean ± SD) and the lighting was adjusted weekly to outdoor conditions, starting with 12:12 h light:dark cycles in late March, which we gradually changed to 14:10 h by the end of April. In April, we collected 50 eggs from each of 12 freshly laid egg strings of the common toad from two ponds in the Pilis-Visegrádi Mountains (Apátkúti tározó: 47.77444° N, 18.98624° E, and Határréti tó: 47.64644° N, 18.90920° E) and one pond in Budapest (Hidegkúti horgásztó: 47.56942° N, 18.95509° E), i.e. four clutches from each population. We housed common toad eggs as described above for agile frogs.

Four days after hatching, when all individuals reached the free-swimming stage (development stage 25; according to Gosner 1960), we started the experiment by haphazardly selecting 36 healthy-looking larvae from each family (36 individuals × 12 families = 432 individuals per species). Tadpoles not used in the experiment were released at the site of their origin. We reared tadpoles individually in opaque plastic containers (18 × 13 × 12 cm) filled with 1 L RSW, arranged in a randomised block design, where each block contained members of one family. Air temperature in the laboratory was 20.1 ± 1.1 °C resulting in 19.0 ± 0.2 °C water temperature in tadpole containers. We changed water in the tadpole rearing containers twice a week and fed tadpoles *ad libitum* with slightly boiled, chopped spinach.

We exposed tadpoles to 19 (unheated control), 28, or 30 °C water temperature for six days, starting 6, 12, or 18 days after hatching (Fig. 1). Thus, thermal treatments were applied during three ontogenetic periods: in early, mid, and late larval stages (hereafter 1^st^, 2^nd^ and 3^rd^ larval period). This resulted in nine treatments with 48 replicates (4 individuals per family × 12 families) in each treatment for each species. In agile frogs, data from the 19 and 30 °C treatments presented here were also used (combined with data from additional treatment groups) for testing another *a-priori* study question, which we published elsewhere (Mikó et al. 2021). We performed thermal treatments in a separate room adjacent to the room where we reared tadpoles. Lighting conditions and room temperature were set to be identical in the two rooms. Immediately before starting thermal treatments, we performed a water change and topped up the RSW to reach a depth of 10 cm (1.7 L RSW in each container during treatment). We placed the containers in 80 × 60 × 12 cm trays filled with tap water to a depth of 8 cm (to avoid floating of the rearing containers), and started to heat the water in the trays to the treatment-specific temperature using thermostated aquarium heaters (Tetra HT 200 in 28 °C treatments and Tetra HT 300 in 30 °C treatments, Tetra GmbH, Melle, Germany). Thereby, water temperature increased gradually to the desired level in ca. two hours, allowing tadpoles to adapt. Opposite to heaters, we placed water pumps (Tetra WP 300) to ensure homogeneous water temperatures, resulting in < 0.5 °C difference among tadpole containers within trays. Overall, this resulted in 28.1 ± 0.4 and 30.0 ± 0.3 °C (mean ± SD) in heated tadpole containers in respective treatments (for details on temperature setting and validation, see the electronic Supporting Information; Fig S1, Table S1). Each tray hosted twelve containers, one from each family (Fig S2), resulting in four trays in each thermal treatment at a time. During the treatment period, we changed water in the tadpole containers every other day with aerated RSW pre-heated to the treatment-specific temperature, and fed tadpoles with a reduced (ca. 1/3) amount of spinach to prevent water fouling and anoxia. Control individuals experienced the same handling and treatment conditions, except that their trays lacked heaters. At the end of the six-day long thermal treatment periods, we changed water with 1 L heated and aerated RSW, removed the containers from the trays and placed them back into their original position in the laboratory, allowing tadpoles to cool down gradually.

**Figure 1:**
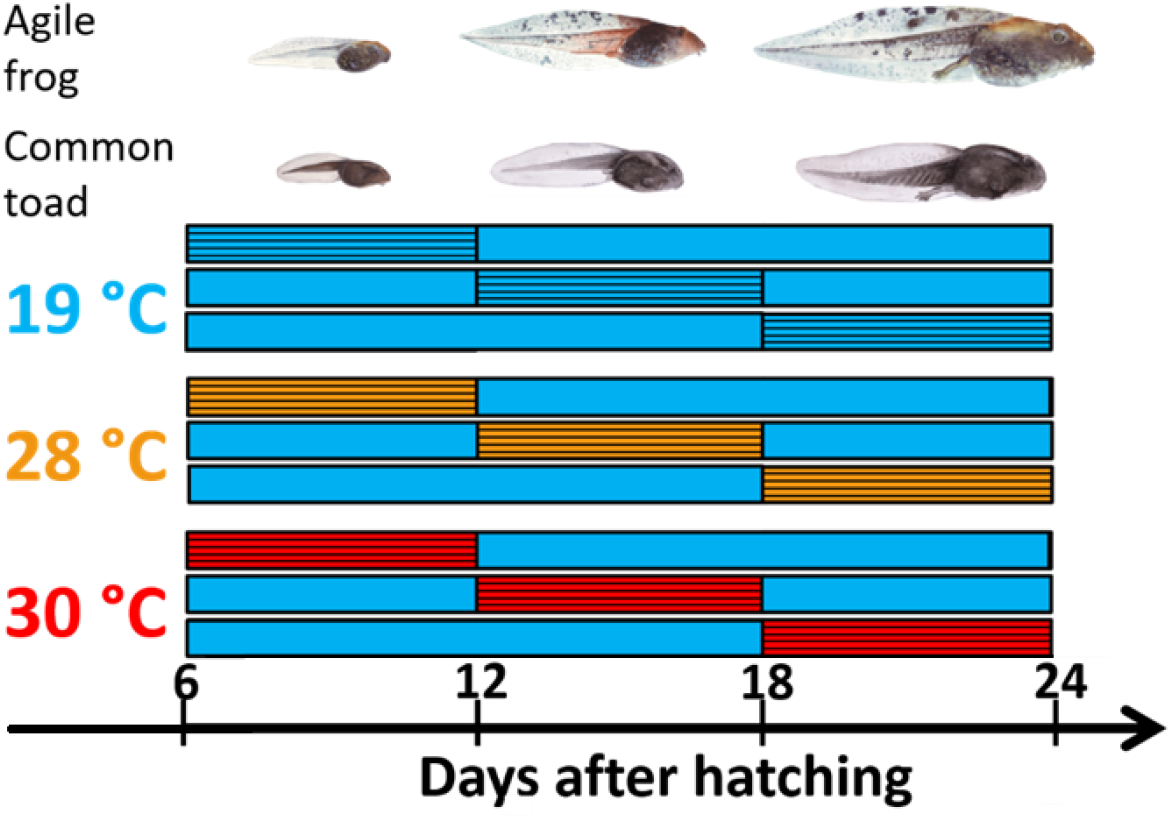
A schematic illustration of experimental treatments. Each horizontal bar represents one treatment group. Striped bars represent periods when tadpoles were exposed to thermal treatments. Orange (28 °C) and red (30 °C) bars symbolize heat treatments, while blue filling represents maintenance at 19 °C. Treatments were identical in both species.

After the last thermal treatments, when tadpoles approached metamorphosis, we checked all rearing containers daily. When an individual started to metamorphose (emergence of forelimbs; development stage 42), we measured its body mass to the nearest 0.1 mg with an analytical balance (Ohaus Pioneer PA-114, Ohaus Europe Gmb, Nanikon, Switzerland), replaced its rearing water with 0.1 L fresh RSW, lifted one side of the container by ca. 2 cm to provide the metamorphs with both water and a dry surface, and covered the container with a transparent, perforated lid. When metamorphosis was completed (complete tail resorption; development stage 46), we placed the individual into a new, lidded container of the same size as before, equipped with wet paper towel lining and a piece of cardboard egg-holder as a shelter. Twice a week, we fed froglets *ad libitum* with small crickets (*Acheta domestica*, instar stage 1-2) sprinkled with a 3:1 mixture of Reptiland 76280 (Trixie Heimtierbedarf GmbH & Co. KG, Tarp, Germany) and Promotor 43 (Laboratorios Calier S.A., Barcelona, Spain) to provide the necessary vitamins, minerals and amino acids. Due to their smaller size, we fed toadlets with springtails (*Folsomia* sp.) in the first three weeks after metamorphosis, and switched to crickets afterwards. For each individual we recorded the dates of starting metamorphosis, completion of tail resorption, and eventual mortality.

Between 6-8 (for agile frogs) or 9-12 (for common toads) weeks after metamorphosis (depending on species and development), when gonads became sufficiently differentiated and easy to observe (Ogielska and Kotusz 2004, Nemesházi et al. 2020), we measured body mass to the nearest 0.01 g and euthanized juvenile individuals in a water bath containing 6.6 g/L tricaine-methanesulfonate (MS-222) buffered to neutral pH with the same amount of Na_2_HPO_4_. We dissected the animals and examined the internal organs under an Olympus SZX12 stereomicroscope (Olympus Europa SE & Co. KG, Hamburg, Germany) at 16× magnification and assigned fat bodies into one of four ordinal categories based on their size: lacking, small, regular-sized, or large. We also categorised phenotypic sex as male (testes), female (ovaries) or uncertain (abnormally looking gonads). Because many animals’ guts contained food remains, we cut out the entire digestive tract, measured its mass to the nearest 0.01 g, and subtracted it from the body mass of juveniles to obtain ‘net body mass’. We removed both feet of euthanized agile frogs and stored them in 96 % ethanol until DNA analyses.

We extracted DNA from agile frog foot samples with Geneaid Genomic DNA Extraction Kit for animal tissue (Thermo Fisher Scientific, Waltham USA) following the manufacturer’s protocol, except that digestion time was 2 hours. We used a recently developed molecular marker set for genetic sexing validated on agile frog populations in Hungary (Nemesházi et al. 2020). We first tested all froglets for the Rds3 marker (≥ 95 % sex linkage) applying high-resolution melting (HRM). We considered an individual to be concordant male or female if its Rds3 genotype was in accordance with its phenotypic sex. Individuals that appeared to be sex-reversed based on the Rds3 marker were also tested using PCR for Rds1 (≥ 89 % sex linkage). For a detailed description of HRM and PCR methods, see Nemesházi et al. (2020). When both markers congruently suggested sex reversal, we considered the given individuals to be sex-reversed. In case of contradiction between the results of analyses based on Rds1 and Rds3, we considered genetic sex to be unknown (Table S2). We did not investigate sex reversal in common toads because phenotypic sex ratios suggested no treatment effects on sex (see Results).

### Statistical analyses

We analysed the data of the two species separately. We assessed treatment effects on survival, length of larval development, body mass at metamorphosis, net body mass at dissection, size of fat bodies, and phenotypic sex ratio. For each dependent variable, we ran a model (see model specifications below) with temperature and treatment period as categorical fixed factors and their interaction, the difference between the mean temperature in each tadpole container and the nominal temperature of the given treatment (measured as described in the electronic Supporting Information) as a numeric covariate, and family nested in population as random factors. We tested the effect of temperature within each treatment period by calculating pre-planned linear contrasts (Ruxton and Beauchamp 2008), correcting the significance threshold for multiple testing using the false discovery rate (FDR) method (Pike 2011). All analyses were conducted in ‘R’ (version 3.6.2), with the ‘emmeans’ package for linear contrasts.

For the analysis of survival, we used Cox’s proportional hazards model (R package ‘coxme’). Individuals were divided into five ordered categories; 1: died during treatment, 2: died after treatment, but before the start of metamorphosis, 3: died during metamorphosis, 4: died after metamorphosis, but before dissection, 5: survived until dissection. Animals that died before the treatment (four agile frog and five common toad larvae) were excluded from survival analyses. We entered the ordinal survival categories as the dependent variable and treated the fifth survival category as censored observations.

To analyse variation in the length of larval development, body mass at metamorphosis and net body mass at dissection, we used linear mixed-effects models (LMM; ‘lme’ function of the ‘nlme’ package), allowing the variances to differ among treatment groups (‘varIdent’ function) because graphical model diagnostics indicated heterogeneous variances. In the analysis of net body mass at dissection, we included age (number of days from finishing metamorphosis to dissection) as a further covariate. In the case of agile frogs, we entered the log-transformed values of the length of larval development to achieve normal distribution of model residuals. For the analysis of fat-body size, we applied cumulative link mixed models (CLMM; ‘clmm’ function of ‘ordinal’ package; Christensen 2015), where we also entered age as a covariate.

To analyse phenotypic sex ratio, first, we excluded those few individuals the gonads of which were not unambiguously categorizable either as male or female (Table S2). Then we analysed the proportion of phenotypic males using phenotypic sex as a binary response variable in generalised linear mixed modelling procedures (GLMM) with binomial error distribution and logit link (‘glmmTMB’ function of the ‘glmmTMB’ package; Brooks et al. 2017). To analyse sex reversal in agile frogs, we could not apply the same modelling framework as for sex ratios because of separation, i.e. sex-reversed individuals were absent in certain treatment groups whereas in some others there was 100% sex reversal. Therefore, we applied six separate analyses comparing the two elevated temperature treatments to their associated controls in each of the three ontogenetic periods using Fisher’s exact tests. The dependent variable was phenotype, i.e. whether or not the individual was sex-reversed. We restricted these analyses to genetic females since heat induces female-to-male sex reversal, and we detected no male-to-female sex reversal. Because of multiple testing, we corrected *P* values using the FDR method.

## Results

Survival of agile frogs that were exposed to 30 °C during either the 1^st^ or the 2^nd^ larval period was significantly reduced (by 56 and 17 %, respectively; Fig. 2, Table 1, Table S3). Survival of common toads also significantly decreased upon exposure to 30 °C (by ca. 33 %), but only if this temperature was applied during the 2^nd^ larval period (Fig.2, Table 2). Thermal treatments that exposed tadpoles to 30 °C in other larval periods (3^rd^ in both species and 1^st^ in common toads) and those involving 28 °C at any period did not affect survival in either species (Table 1-2).

**Table 1:**
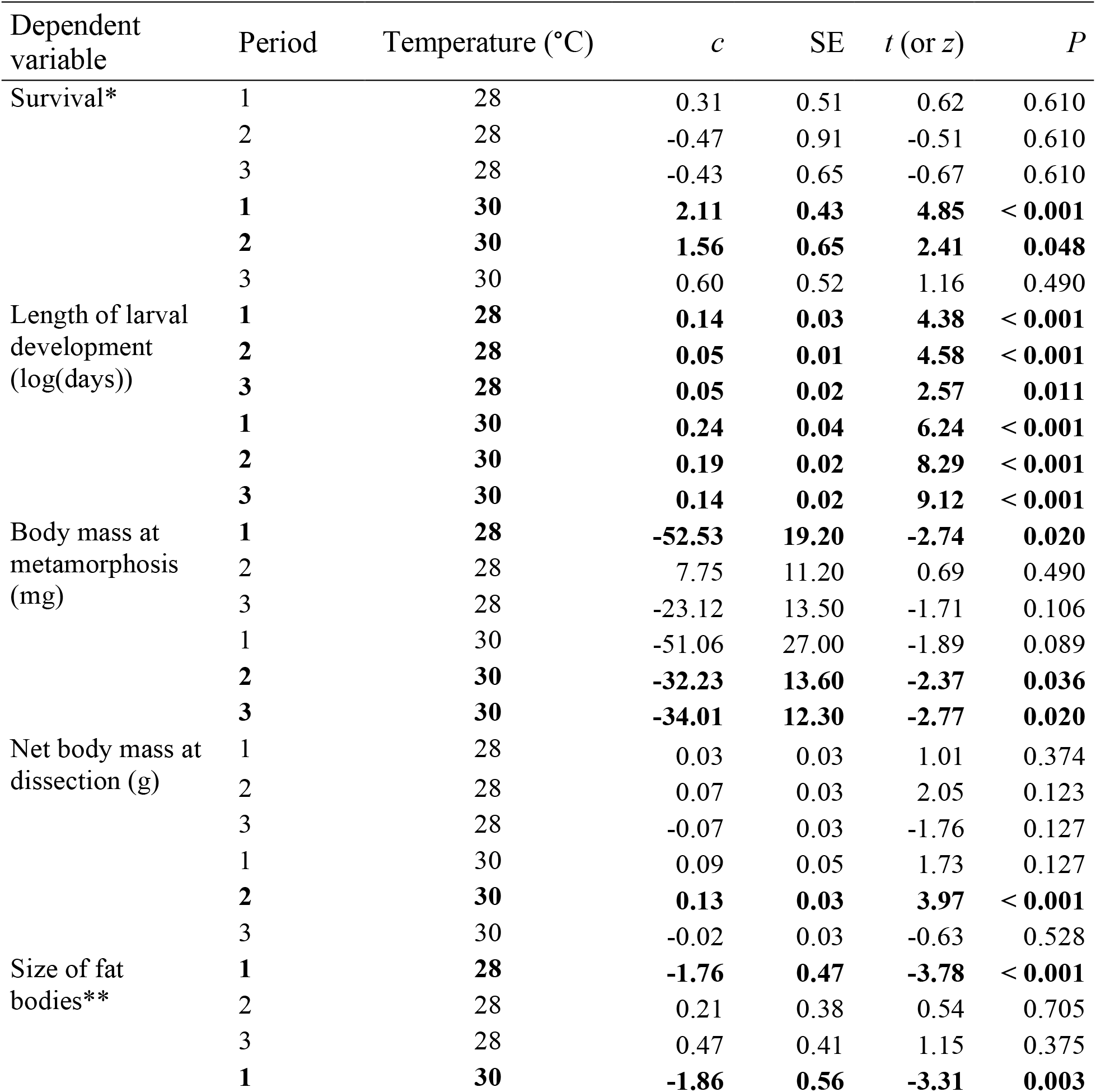
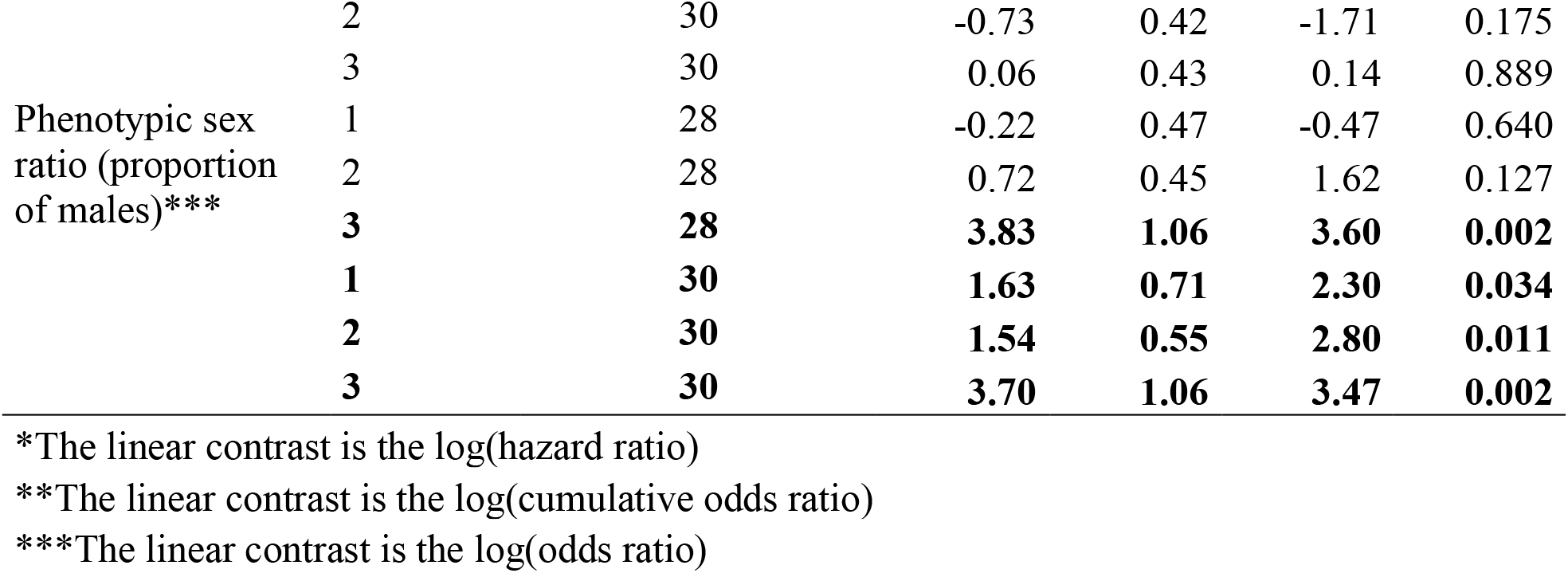
Agile frog responses to heat by the timing of exposure (1^st^, 2^nd^ and 3^rd^ larval period) and the applied temperature. Results represent pre-planned comparisons from the models shown in Table S2, comparing each period and temperature combination to the 19°C treatment in the corresponding period. Linear contrasts (*c*), associated standard errors (SE), t-values (z-values in case of Cox’s proportional hazards model in the analyses of survival) and *P*-values adjusted using the FDR method are reported. Treatment groups that differed significantly (P < 0.05) from their corresponding controls are highlighted in bold.

**Table 2:**
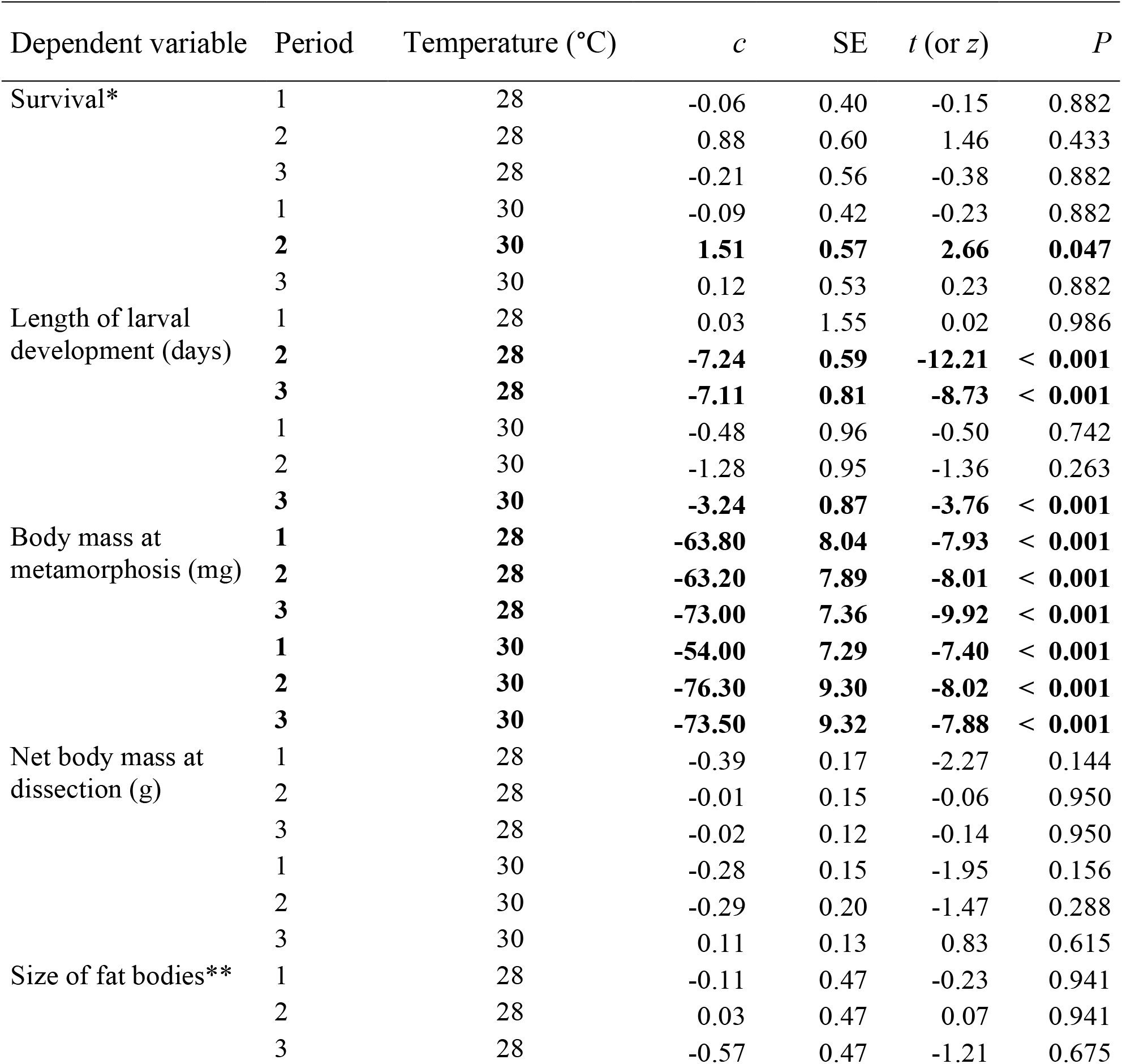
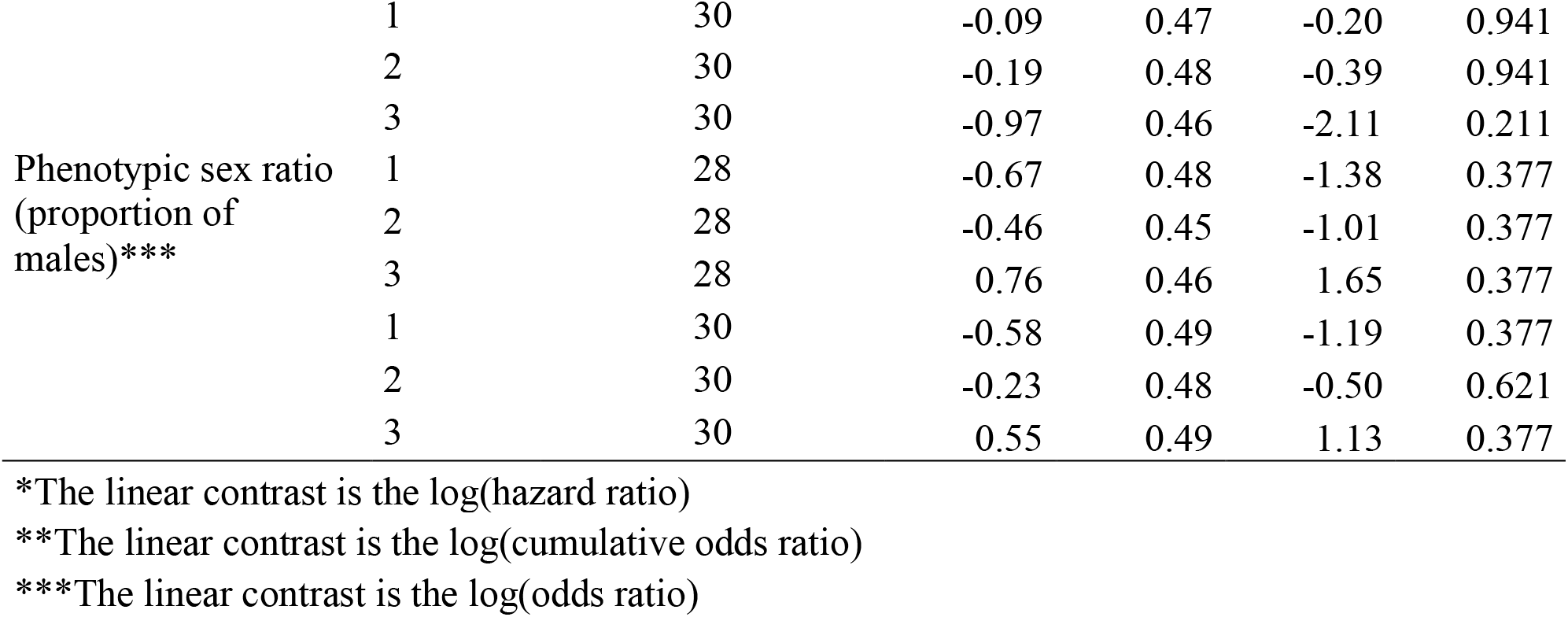
Common toad responses to heat by the timing of exposure (1^st^, 2^nd^ and 3^rd^ larval period) and the applied temperature. Results represent pre-planned comparisons from the models shown in Table S2, comparing each period and temperature combination to the 19°C treatment in the corresponding period. Linear contrasts (*c*), associated standard errors (SE), t-values (z-values in case of Cox’s proportional hazards model in the analyses of survival) and *P*-values adjusted using the FDR method are reported. Treatment groups that differed significantly (P < 0.05) from their corresponding controls are highlighted in bold.

**Figure 2:**
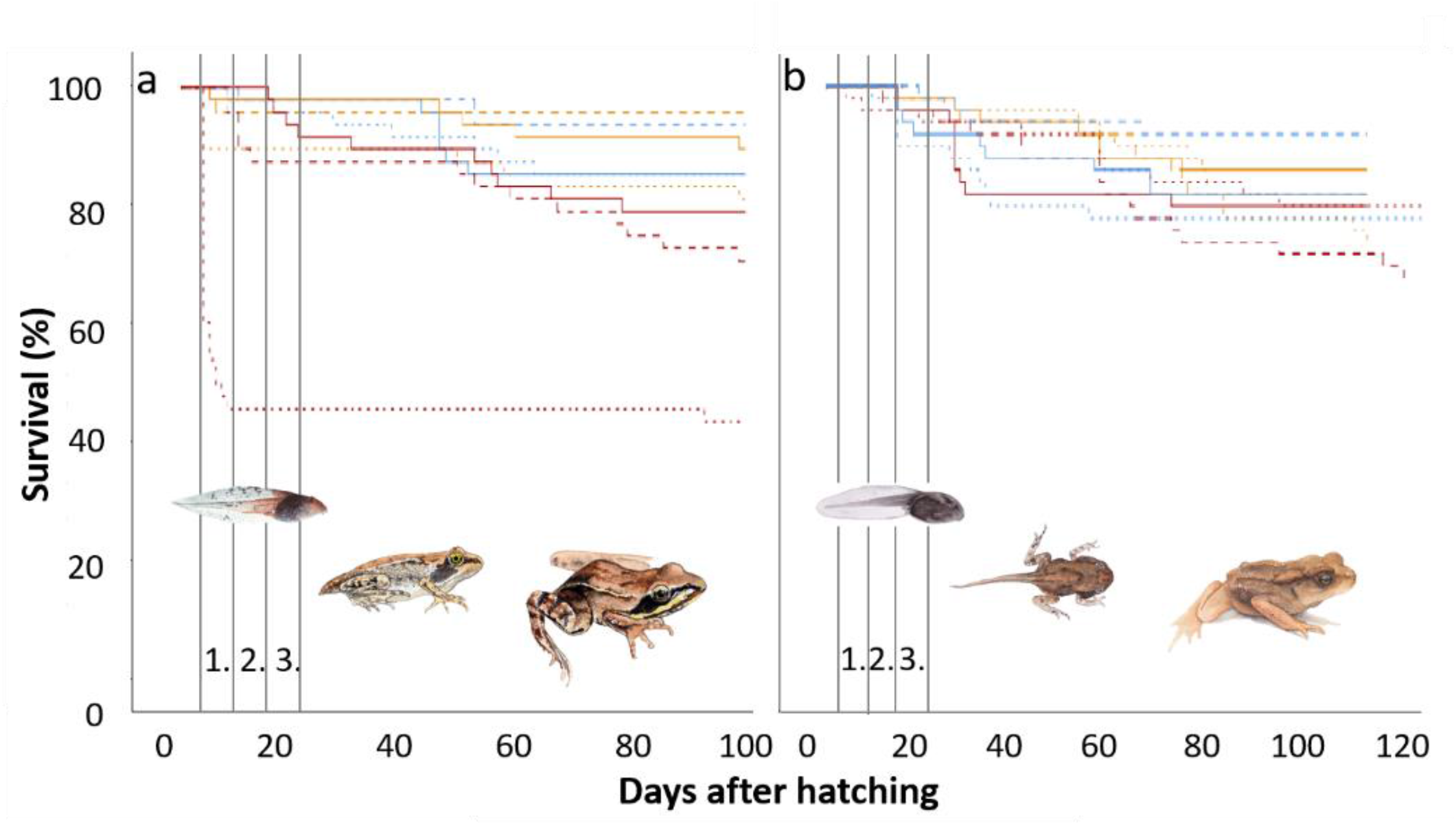
Survival of agile frogs (a) and common toads (b) during the experiment over time in each treatment group. Blue lines represent controls maintained at 19 °C throughout, orange lines represent treatment groups exposed to 28 °C, red lines represent treatment groups exposed to 30 °C; dotted lines represent individuals exposed to thermal treatments during the 1^st^ larval period, dashed lines those exposed during the 2^nd^ larval period, and solid lines those exposed during the 3^rd^ larval period. Numbered vertical lanes depict the respective larval periods when thermal treatments were performed. Note that the experiment lasted longer for common toads than for agile frogs.

Length of larval development of agile frogs was significantly prolonged by all thermal treatments applied in all larval periods (Fig. 3, Table 1 and S3). By contrast, in common toads, the length of larval development was not affected when tadpoles were exposed to 28 °C during the 1^st^ larval period, but larvae that were exposed to this temperature during the 2^nd^ and 3^rd^ larval period developed faster compared to their control groups (Fig. 4, Table 2 and S3). When common toad tadpoles were exposed to 30 °C, their larval development was only shortened upon exposure during the 3^rd^ larval period but remained unaffected if treated in the 1^st^ or 2^nd^ larval period (Fig 4, Table 2 and S3).

**Figure 3:**
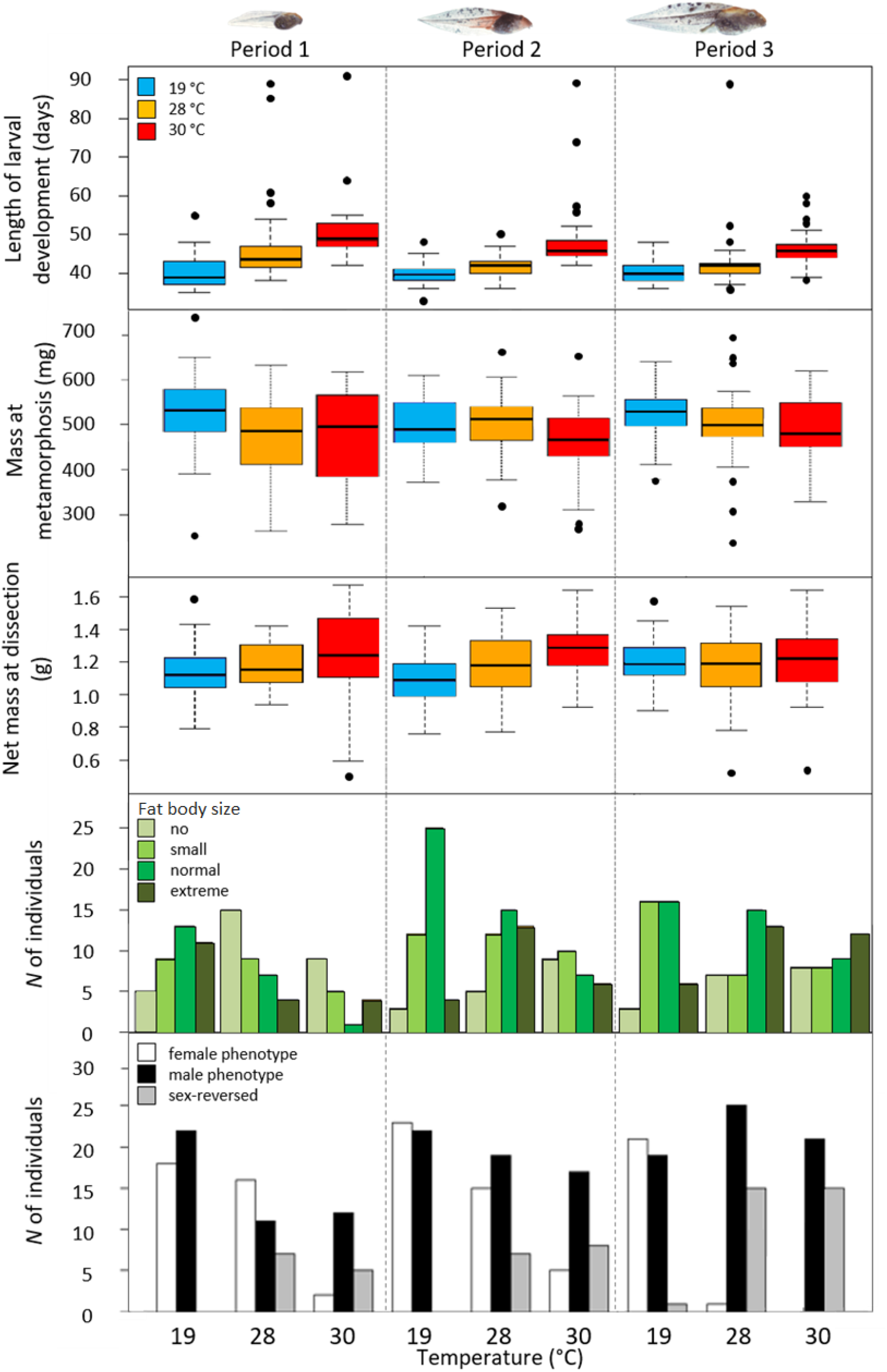
Agile frog responses to thermal treatments in terms of the length of larval development, body mass at metamorphosis, net body mass at dissection, the size of fat bodies and phenotypic sex ratios in juveniles. In boxplots, horizontal lines and boxes represent medians and interquartile ranges (IQR), respectively, while whiskers extend to IQR ± 1.5×IQR and dots indicate more extreme data points.

**Figure 4:**
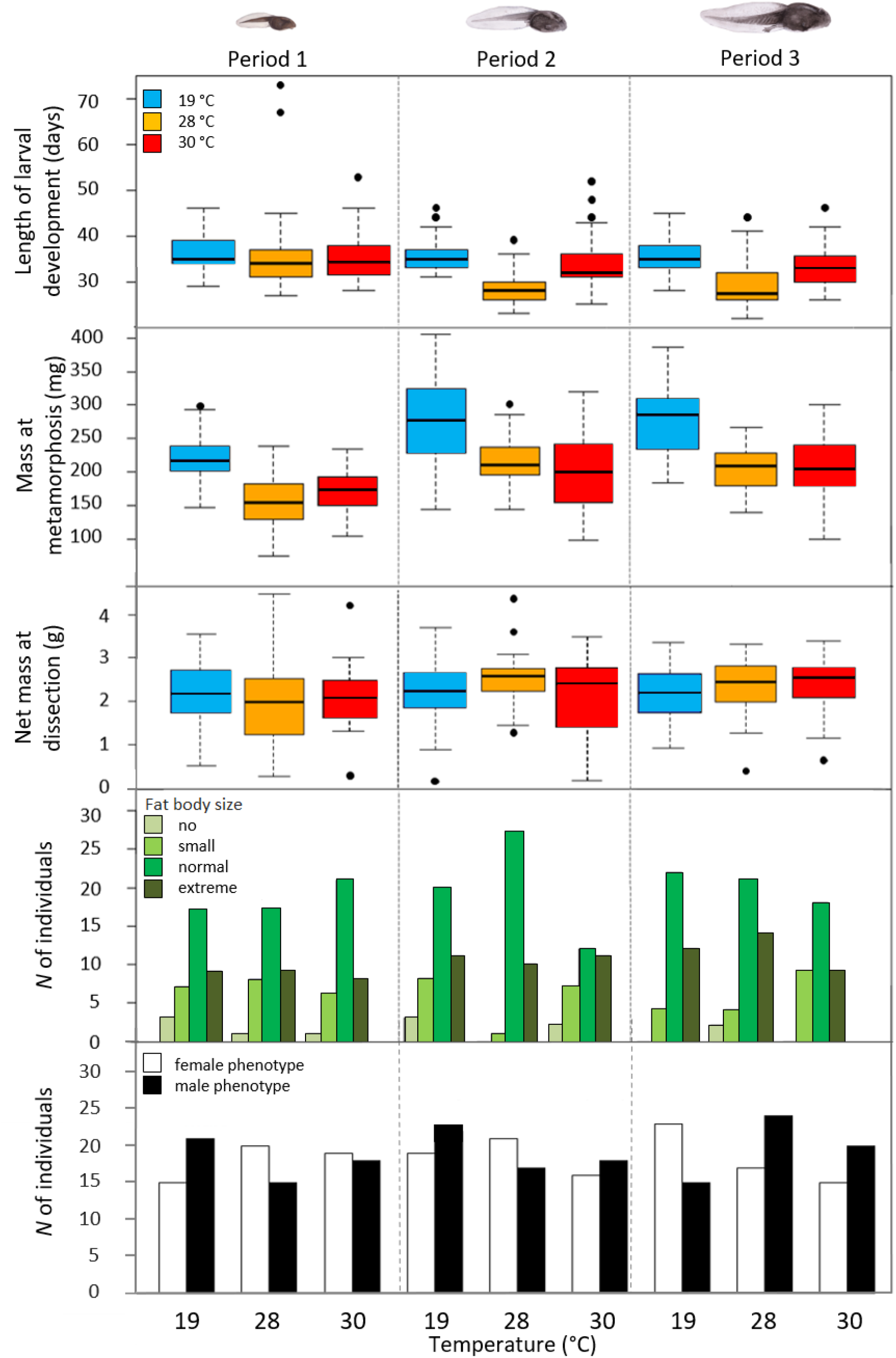
Common toad responses to thermal treatments in terms of the length of larval development, body mass at metamorphosis, net body mass at dissection, the size of fat bodies and phenotypic sex ratios in juveniles. In boxplots, horizontal lines and boxes represent medians and interquartile ranges (IQR), respectively, while whiskers extend to IQR ± 1.5×IQR and dots indicate more extreme data points.

Body mass at metamorphosis was significantly reduced in agile frogs by the 28 °C thermal treatment if applied during the 1^st^ larval period but was not affected if 28 °C was applied later on (Fig 3, Table 1). Exposure to 30 °C tended to decrease body mass at metamorphosis when applied in the 1^st^ larval period and exerted a significant negative effect during the 2^nd^ and 3^rd^ larval period (Fig 3, Table 1 and S3). In common toads, both thermal treatments applied in all larval periods resulted in significantly reduced body mass at metamorphosis (Fig 4, Table 2 and S3).

At dissection, net body mass of juvenile agile frogs was only increased in animals treated with 30 °C during the 2^nd^ larval period, but remained unaffected in all other treatment groups (Fig 3, Table 1 and S3). Thermal treatments applied in any larval period did not affect the net body mass of common toads (Fig 4, Table 2 and S3). The number of days between metamorphosis and dissection positively affected net body mass at dissection in both species (Table S3).

The size of fat bodies was significantly smaller in juvenile agile frogs as a result of both thermal treatments, but only upon exposure during the 1^st^ larval period and not during later periods (Fig 3, Table1). In juveniles of the common toad the size of fat bodies was unaffected by thermal treatments applied in any larval period (Fig 4, Table 2), and positively correlated with the age of juveniles (Table S3).

Phenotypic sex ratio in agile frogs was affected by exposure to elevated temperature: exposure to 28 °C during the 3^rd^ larval period (but not in the earlier periods) caused a significant shift towards a male-biased sex ratio, and treatment with 30 °C in all larval periods resulted in highly male-biased sex ratios (Fig 3, Table 1, S2 and S3). Accordingly, the proportion of agile frog individuals that underwent heat-induced sex reversal was significantly higher (between 30 and 100 % of genetic females) at both temperatures and in all treatment periods compared to the respective control groups (≤ 4.5 %, all *P* ≤ 0.012; Fig 3, Table S2). In contrast, none of the thermal treatments applied in either larval period had any effect on the phenotypic sex ratio of juvenile common toads (Fig 4, Table 2, S2 and S3).

## Discussion

Our results demonstrate that high temperatures experienced for six days during larval development can negatively affect the survival, growth, somatic and sexual development of amphibians, but the severity of these effects depends on the intensity and timing of thermal stress and can largely differ between species. Agile frogs proved to be more sensitive: in this species, all studied variables were affected by one or more heat treatments, and almost all of the resulting changes are likely disadvantageous for individual fitness and population viability (Fig. 5). In contrast, for common toads, the only consistent effect of thermal stress was reduced mass at metamorphosis and, in a few treatments, faster larval development, while we observed barely any effect on survival and no lasting developmental effects in juveniles (Fig. 5). These results highlight that even sympatric species that are relatively similar in their ecology may be affected very differently by heat waves.

**Figure 5:**
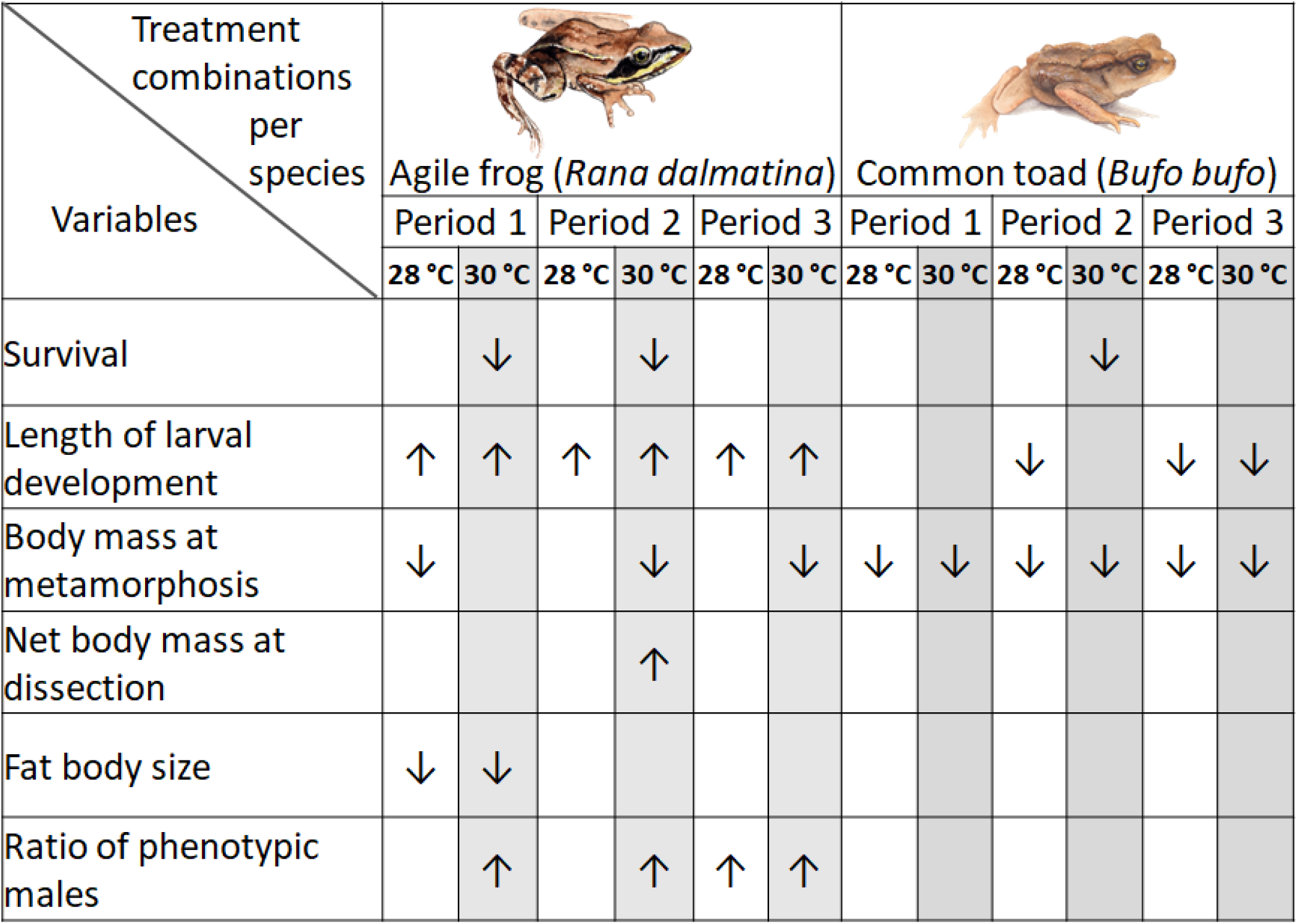
Summary of responses by the two species to the simulated heat waves in different larval periods. Arrows show the direction of the observed change in the given variable relative to the respective control group, not the advantageousness or harmfulness of the effect. Separation is aided by different colours.

Survival rate in both species was decreased by exposure to 30 °C, but only if tadpoles experienced it relatively early on during their development (during 1^st^ and 2^nd^ larval periods). Temperatures of around 30 °C throughout the entire larval development often resulted in decreased survival in earlier studies (Bellakhal et al. 2014, Goldstein et al. 2017, Phuge 2017, Lambert et al. 2018). Our results suggest that the adverse effect of elevated temperature on larval survival depends on the species and on the timing of exposure, indicating a peak in thermosenitivity during the early stages of larval development (in addition to the increased thermosensitivity of the final larval stages, directly before the onset of metamorphosis (Floyd 1983), which we did not study). This is in line with many previous studies suggesting that the earliest life stages of amphibians are the most susceptible to several stress factors such as chemicals, parasites, poor environmental conditions and pesticides (Ortiz-Santaliestra et al. 2006, Holland et al. 2007, Crespi and Warne 2013, Mikó et al. 2017). The energetically costly cellular repairing mechanisms and the maintenance and restoration of homeostasis during and after thermal stress compromise higher-level functions that are necessary for survival (Williams et al. 2016). Furthermore, dissolved oxygen level in the water decreases with rising temperature (Stefan et al. 2001, Fang and Stefan 2009), which in turn can cause hypoxia and oxidative stress in tadpoles (Lushchak 2011, Freitas and Almeida 2016). High temperature may also accelerate bacterial bloom in the water (Ferreira and Chauvet 2011), potentiating the accumulation of opportunistic pathogens. All of these processes might contribute to mortality observed in experiments involving thermal treatments and, under natural conditions, during or after heat waves.

Timing of metamorphosis and body mass at metamorphosis are crucial components of fitness in amphibians. Earlier metamorphosis allows for leaving the more hazardous aquatic environment faster (Denver 1997), and allows for a longer post-metamorphic growth period compared to late-metamorphosing individuals, which in turn leads to increased survival during the first hibernation (Altwegg and Reyer 2003, Üveges et al. 2016). In the present study, the simulated heat waves prolonged larval development in agile frogs but shortened it (when heat was experienced in the late larval period) in common toads, whereas mass at metamorphosis decreased after heat exposure in both species (although in agile frogs the latter effect was only significant in a few treatment groups). According to the temperature-size rule (Kozłowski et al. 2004), high temperatures are associated with increased metabolic rates and accelerated development in larval anurans (Álvarez and Nicieza 2002, McLeod et al. 2013, Courtney Jones et al. 2015), which results in earlier metamorphosis at a smaller body size (Laugen et al. 2003, Niehaus et al. 2006). Our results likely documented this relationship between development and growth in common toads. However, in agile frogs, this relationship was disrupted by heat treatments, most probably because the applied temperatures acted as severe stressors. This result aligns with the observation that larvae of the common toad are more thermophilic than those of agile frogs, as suggested by a higher critical thermal maximum and higher preferred temperatures in the former than in the latter (Hettyey, personal communication).

Stress experienced early in life can have long-lasting consequences, such as small adult size and limited energy reserves (Crespi and Warne 2013, Jonsson and Jonsson 2014). However, in our study, the reduced mass at metamorphosis in heat-treated groups did not persist into juvenility: after a few months of post-metamorphic growth, we found no differences in body mass or fat reserves in either species. There were only two exceptions to this: in juvenile agile frogs, fat bodies were smaller if they received either heat treatment in the 1^st^ larval period, and unexpectedly, their body mass was larger after exposure to 30 °C applied during the 2^nd^ larval period. The death of lighter individuals likely contributed to the equalization of juvenile body mass among treatment groups, given that most individuals that died between the onset of metamorphosis and dissection had a lower body mass at metamorphosis than conspecifics that survived until the end of the experiment in both species (Welch’s tests; agile frogs: *t* = -3.54, *df* = 32.0, *P* = 0.001, common toads: *t* = -9.30, *df* = 53.9, *P* < 0.001). A further contributing factor may be compensatory growth (Squires et al. 2010, Hector et al. 2012). Nonetheless, compensatory growth can have hidden costs (Stoks et al. 2006, De Block and Stoks 2008, Murillo-Rincón et al. 2017), so that the lack of among-treatment differences in juvenile mass does not necessarily indicate the absence of long-term malign consequences of high temperatures experienced during larval life. Indeed, the majority of juvenile agile frogs completely lacked fat bodies if they were exposed to heat during the 1^st^ larval period. Fat bodies in amphibians are major energy stores that are vital to survival (Scott et al. 2007) and regulate processes related to reproduction (Pierantoni et al. 1983, Girish and Saidapur 2000). Consequently, high temperatures experienced during early ontogeny may have long-lasting negative effects on the survival and reproductive potential of agile frogs, which may compromise population persistence. The observation that the size of fat bodies was not affected by thermal treatments in common toads confirms that these are more tolerant to high temperatures than agile frogs, and, more generally, reinforces the hypothesis that there is large among-species variation also in the long-term consequences of thermal stress.

Sex reversal can occur naturally in wild populations of agile frogs (Nemesházi et al. 2020) and other species (Alho et al. 2010, Lambert et al. 2019, Xu et al. 2021), but high temperature can increase its frequency in a wide range of ectothermic vertebrates (Baroiller and D’Cotta 2016, Ruiz-Garciá et al. 2021, Whiteley et al. 2021). In our study, six-day 30 °C heat waves caused male-biased sex ratios via sex reversal in agile frogs, and the same effect was induced by exposure to 28 °C in the 3^rd^ larval period. These results align with previous studies documenting altered sex ratios in several anuran species where larvae were raised at high temperatures throughout their development (Ruiz-Garciá et al. 2021), and additionally suggest that the sensitivity of sex determination to elevated temperature increases close to the end of larval development. Our findings caution that heat waves lasting for only a few days during tadpole development can trigger sex reversal, which may have wide-ranging consequences including skewed sex ratios and lowered population viability (Bókony et al. 2017, Wedekind 2017, Nemesházi et al. 2021). However, our observation that the same thermal treatments did not affect phenotypic sex ratios in common toads suggests that there is considerable interspecific variation in the thermosensitivity of sexual development.

Heat treatment is a promising mitigation method against amphibian chytridiomycosis (Chatfield and Richards-Zawacki 2011, Geiger et al. 2011, Hettyey et al. 2019). Our results, however, underline the importance of pre-assessing the thermal sensitivity of each species, including that of their sexual development. Based on our results, thermal treatment at 30 °C could be applied for six days to common toads, which would likely lead to *Bd* clearance, or at least to a drastic suppression of *Bd* growth (Retallick and Miera 2007, Chatfield and Richards-Zawacki 2011, Geiger et al. 2011). This treatment could be recommended in specific situations, such as epizootic outbreaks, when the benefits clearly outweigh the costs arising from decreased body mass at metamorphosis, or when the latter can be compensated for (e.g. by supplemental feeding). In agile frogs, treatment with 28 °C during the 2^nd^ larval period (days 12-18 after hatching) was the only treatment combination without adverse effects on most life-history traits and sexual development. Although this treatment also caused somewhat lengthened larval development, this cost may be negligible (especially so in captivity) considering the benefit of Bd clearance. Whether treatment with temperatures lower than 28 °C would be applicable without costs and still suppresses *Bd* growth sufficiently needs further investigation (Hettyey et al. 2019). A further possibility to explore is that under controlled conditions, capitalising on the feminizing effect of estrogens or other estrogenic chemicals might make thermal treatment of *Bd*-infected animals potentially suitable also for species with thermally sensitive sex determination (Kitano et al. 2012).

In conclusion, our study demonstrates that species can differ in a multitude of ways in how they are affected by short periods of elevated temperatures which are similar in magnitude to those occurring in natural water bodies during heat waves. Most importantly, we demonstrate that already 28 °C can have surprisingly severe consequences for larvae of a thermosensitive anuran, where the strength of effects depends largely on the developmental stage of individuals that become exposed to the heat. At the same time, even 30 °C experienced any time during larval development does little harm to individuals of another sympatric species. Such species-specific differences should be examined in a wide range of taxa and considered when evaluating the impact of climate change on amphibians, and also in the development of mitigating methods against chytridiomycosis.

## Acknowledgements

We are thankful to Zsófia Boros, Dóra Holly, Boglárka Jaloveczki, Csenge Kalina, Eszter Nádai-Szabó, Stephanie Orf and Gergely Tarján for their help during the experiment and data archiving. We thank Gergő Tholt and the NÖVI Department of Zoology for providing us with their stereomicroscope and camera. Bálint Bombay made the paintings of tadpoles and juvenile frogs. The study was funded by the Lendület Programme of the Hungarian Academy of Sciences (MTA, LP2012-24/2012), an FP7 Marie Curie Career Integration Grant (PCIG13-GA-2013-631722) and the National Research, Development and Innovation Office of Hungary (NKFIH, grants 115402 and 135016 to V.B., 124708 to O.I.H., and 124375 to A.H.). The authors were supported by the János Bolyai Research Scholarship of the Hungarian Academy of Sciences (to V.B., A.H., and O.I.H.), the New National Excellence Program of the Ministry for Innovation and Technology from the source of the National Research, Development and Innovation Fund (ÚNKP-20-5 and ÚNKP-21-5 to V.B. and A.H., ÚNKP-19-4 to A.H., ÚNKP-21-4 to J.U., and ÚNKP-19-3, ÚNKP-20-3, ÚNKP-21-3 to A.K.), the Ministry of Human Capacities (National Program for Talent of Hungary, NTP-NFTÖ-18-B-0412 to V.V., NTP-NFTÖ-17-B-0317 to E.N.), and the Austrian Agency for International Cooperation in Education & Research (OeAD-GmbH; ICM-2019-13228 to E.N.). N.U. and D.H. were supported by the Young Investigators Programme of the Hungarian Academy of Sciences. Experimental procedures were approved by the Ethical Commission of the Plant Protection Institute, and permissions were issued by the Government Agency of Pest County (PE/KTF/3596-6/2016, PE/KTF/3596-7/2016 and PE/KTF/3596-8/2016). The experiments were carried out according to recommendations of the EC Directive 86/609/EEC for animal experiments (http://europa.eu.int/scadplus/leg/en/s23000.htm).

The authors have no conflict of interest to declare.

## Data availability statement

The data that support the findings of this study are openly available in figshare at http://doi.org/10.6084/m9.figshare.17197847.

## Supporting Information to

### Measurements validating temperature in heat treatments

We validated the heating setup by repeatedly measuring water temperature (± 0.1 °C) in tadpole containers of each position in each tray, as well as water temperature in the trays in which treatments took place. Before the experiment, we measured these temperatures ten times on two consecutive days with a Greisinger digital thermometer (GTH175/PT). After termination of the experiment, we repeated these measurements five times. To detect eventual temperature fluctuations during each treatment, twice per day we checked water temperature in all trays using the digital thermometer. Furthermore, data loggers (Onset HOBO Pendant Temperature/Light 8K Data Logger; one per each tray) recorded temperature in the trays every 30 minutes during the treatments. We did not measure temperature in the tadpole containers during the treatment periods in order to avoid stress and injury as a result of stirring the water. We did not detect considerable temperature fluctuations during the treatments (Fig S1, Table S1), and temperature readings were very similar before and after the experiment in each container position. Temperature did vary somewhat among containers in different positions within trays (maximal temperature difference within a tray at 19 °C: 1.3 °C; at 28 °C: 1.5 °C; at 30 °C: 1.5 °C), but this variation was highly consistent over time. We calculated the difference between the actual (experienced by the tadpoles) and nominal temperature of the given treatment for each tadpole container by subtracting the nominal temperature from the mean water temperature (measured before and after the treatments in each container with the digital thermometer). This method minimised the disturbance caused to animals during the experiment while delivering accurate data on the temperatures experienced by the tadpoles.

**Figure S1.**
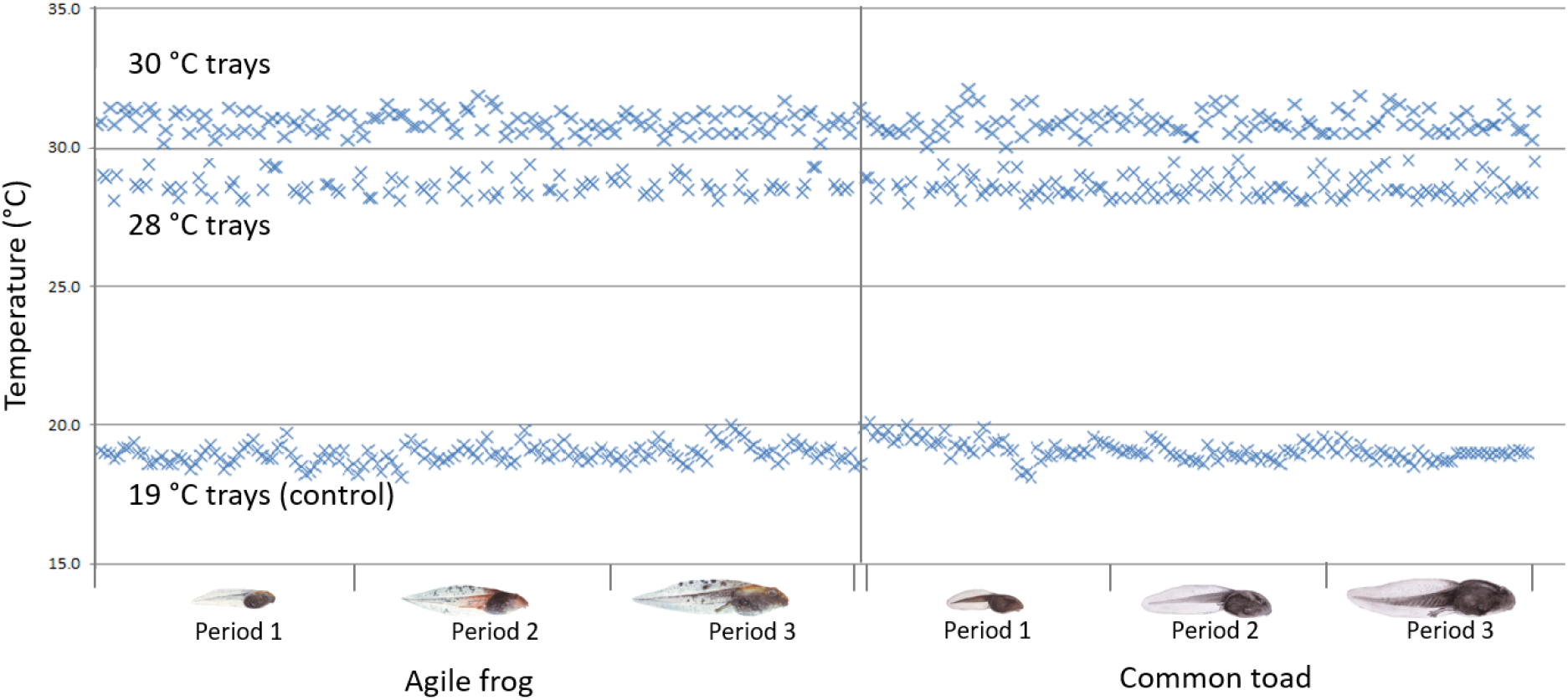
Water temperatures in the trays during treatment periods show minimal temperature fluctuations. Note that the water temperature in the heated trays was always warmer than the temperature in the tadpoles’ containers (set to be as close to the nominal temperature as possible; Table S1).

**Table S1.** Minimal (T_min_), maximal (T_max_) and mean (T_mean_) temperatures in the heated trays during temperature treatments. Mean diff. represents the average difference in water temperature between the tadpoles’ containers and the trays, since water temperature in the heated trays was always warmer than the temperature in the tadpoles’ containers.

| Tray | $T_{\min}$ (°C) | $T_{\max}$ (°C) | $T_{\text{mean}}$ (°C) | Mean diff. (± SE) |
| --- | --- | --- | --- | --- |
| 30°C /1 | 30.6 | 31.6 | 31.0 | 1.1 (± 0.13) |
| 30°C /2 | 30.2 | 31.6 | 30.8 | 1.1 (± 0.10) |
| 30°C /3 | 30.4 | 31.6 | 30.9 | 0.5 (± 0.09) |
| 30°C /4 | 30.5 | 31.3 | 30.8 | 0.7 (± 0.09) |
| 28°C /1 | 28.2 | 28.9 | 28.5 | 0.4 (± 0.06) |
| 28°C /2 | 28.4 | 29.4 | 28.8 | 0.1 (± 0.07) |
| 28°C /3 | 28.0 | 29.5 | 28.5 | 0.5 (± 0.06) |
| 28°C /4 | 28.1 | 29.5 | 28.6 | 1.1 (± 0.08) |

**Figure S2.**
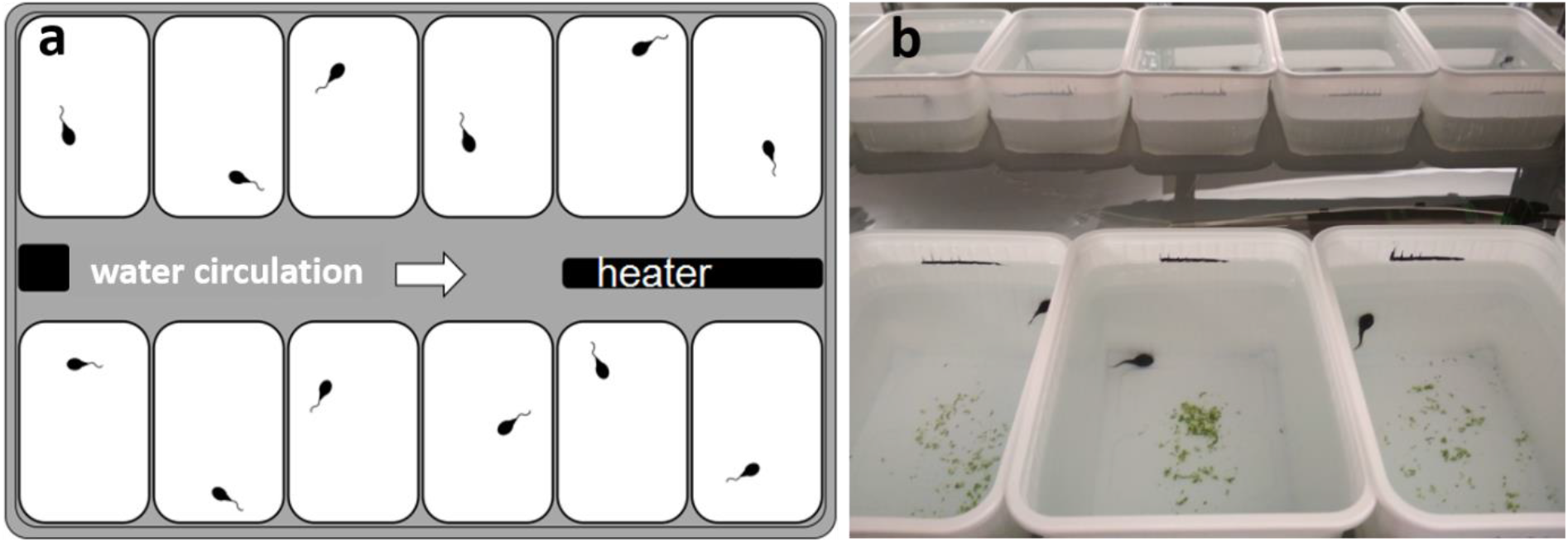
Schematic representation (a) and *in situ* photograph (b) of the heating system used in the thermal treatments.

**Table S2.** Phenotypic sex ratio (% males) in each treatment group. In case of agile frogs, genetic sex ratio and female-to-male sex-reversal rate (% of phenotypic males in genetic females) are also shown.

| Species | Treatment group | Dissected (N) | Phenotypic sex ratio (% male)<sup>†</sup> | Genetic sex ratio (% male)<sup>‡</sup> | Female-to-male sex reversal (%) |
| --- | --- | --- | --- | --- | --- |
| Agile frog | Period 1 |  |  |  |  |
|  | Control | 40 | 55.0 | 55.0 | 0.0 |
|  | 28 °C | 39 | 52.9 | 32.4 | 30.4 |
|  | 30 °C | 19 | 89.5* | 63.2 | 71.4 |
|  | Period 2 |  |  |  |  |
|  | Control | 45 | 48.9 | 48.9 | 0.0 |
|  | 28 °C | 46 | 63.4 | 46.3 | 31.8 |
|  | 30 °C | 32 | 83.3* | 56.6 | 61.5 |
|  | Period 3 |  |  |  |  |
|  | Control | 41 | 48.8 | 46.3 | 4.5 |
|  | 28 °C | 43 | 97.5* | 60.9 | 93.8 |
|  | 30 °C | 37 | 100.0* | 58.3 | 100.0 |
| Common toad | Period 1 |  |  |  |  |
|  | Control | 36 | 58.3 |  |  |
|  | 28 °C | 35 | 42.9 |  |  |
|  | 30 °C | 37 | 48.6 |  |  |
|  | Period 2 |  |  |  |  |
|  | Control | 42 | 54.8 |  |  |
|  | 28 °C | 39 | 43.6 |  |  |
|  | 30 °C | 34 | 52.9 |  |  |
|  | Period 3 |  |  |  |  |
|  | Control | 38 | 39.5 |  |  |
|  | 28 °C | 41 | 58.5 |  |  |
|  | 30 °C | 36 | 55.6 |  |  |

\*Sex ratios that differ significantly from 1:1 according to Fisher's exact tests
†Excluding those individuals whose gonads were not unambiguously categorizable either as male or female (3 agile frogs: 2 individuals at 28 °C in the 2<sup>nd</sup> larval period and 1 individual at 30 °C in the 2<sup>nd</sup> larval period; 2 common toads: 1 individual at 28 °C in the 2<sup>nd</sup> larval period and 1 individual at 30 °C in the 3<sup>rd</sup> larval period)
‡Excluding those individuals whose genetic sex was unknown due to contradiction between the Rds1 and Rds3 markers' results (at 28 °C 1 individual in the 1<sup>st</sup>, and 2 individuals in the 3<sup>rd</sup> larval period; at 30 °C 1 individual each in the 2<sup>nd</sup> and 3<sup>rd</sup> larval periods)

**Table S3.**
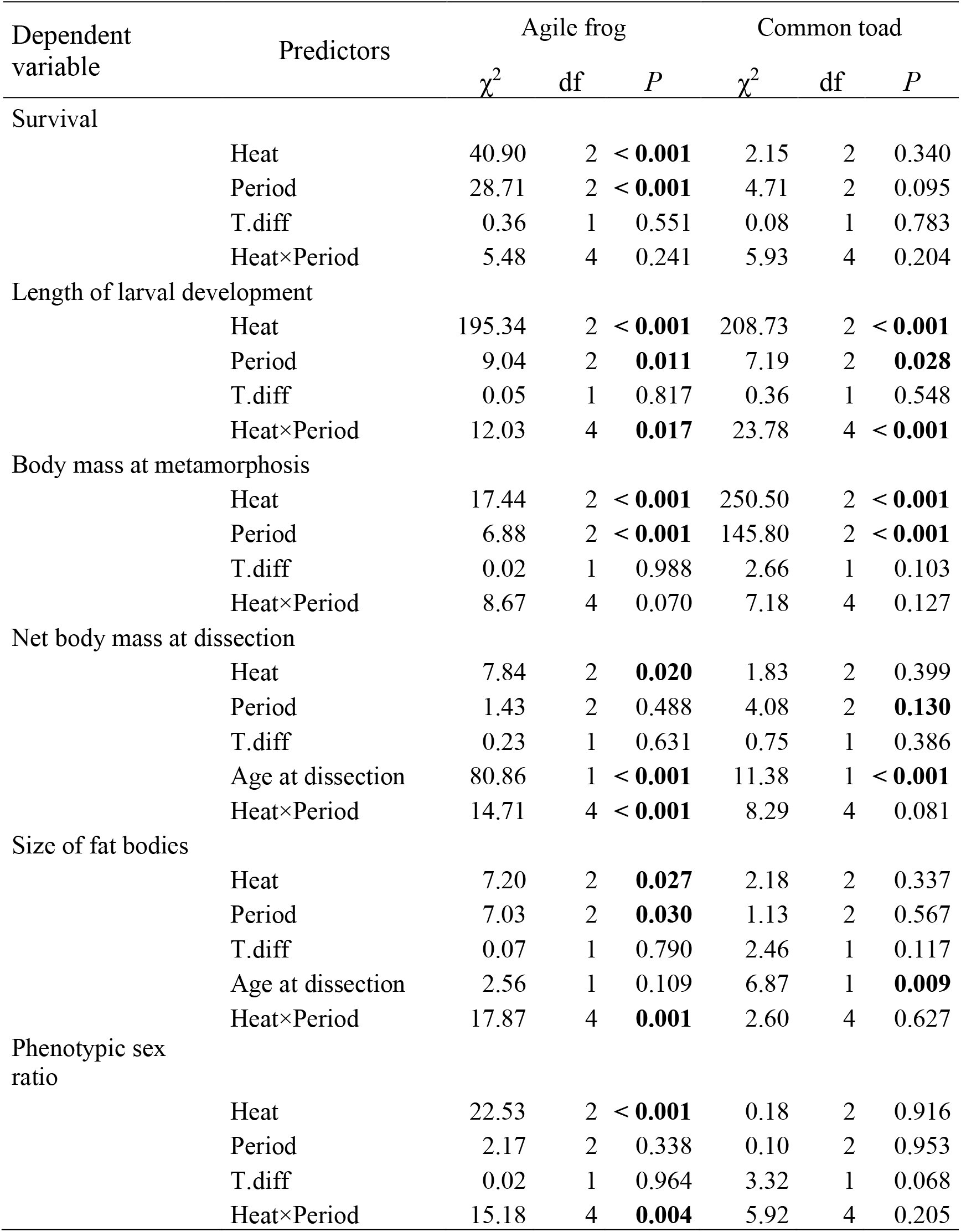
Type-2 analysis-of-deviance tables of the statistical models. Significant effects (*P* < 0.05) are highlighted in bold. The covariate “T.diff” is the difference between mean temperatures in each tadpole container and the nominal temperature of the given treatment.

## References

Alho, J. S. et al. 2010. Sex reversal and primary sex ratios in the common frog (Rana temporaria). - Mol. Ecol. 19: 1763–1773.

Altwegg, R. and Reyer, H.-U. 2003. Patterns of natural selection on size at metamorphosis in water frogs. - Evolution (N. Y). 57: 872–882.

Álvarez, D. and Nicieza, A. G. 2002. Effects of temperature and food quality on anuran larval growth and metamorphosis. - Funct. Ecol. 16: 640–648.

APHA et al. 1992. Standard methods for the examination of water and wastewater - American Public Health Association, American Water Works Association, Water Environment Federation. - American Public Health Association.

Arnfield, A. J. 2003. Two decades of urban climate research: A review of turbulence, exchanges of energy and water, and the urban heat island. - Int. J. Climatol. 23: 1–26.

Baroiller, J. F. and D’Cotta, H. 2016. The reversible sex of gonochoristic fish: Insights and consequences. - Sex. Dev. 10: 242–266.

Becker, C. G. et al. 2012. Disease Risk in temperate amphibian populations is higher at closed-canopy sites. - PLoS One 7: e48205.

Bellakhal, M. et al. 2014. Effects of temperature, density and food quality on larval growth and metamorphosis in the north African green frog Pelophylax saharicus. - J. Therm. Biol. 45: 81–86.

Berger, L. et al. 1998. Chytridiomycosis causes amphibian mortality associated with population declines in the rain forests of Australia and Central America. - Proc. Natl. Acad. Sci. U. S. A. 95: 9031–9036.

Berger, L. et al. 2004. Effect of season and temperature on mortality in amphibians due to chytridiomycosis. - Aust. Vet. J. 82: 31–36.

Blaustein, A. R. et al. 2001. Amphibian breeding and climate change. - Conserv. Biol. 15: 1804–1809.

Blaustein, A. R. et al. 2005. Interspecific variation in susceptibility of frog tadpoles to the pathogenic fungus Batrachochytrium dendrobatidis. - Conserv. Biol. 19: 1460–1468.

Blaustein, A. R. et al. 2010. Direct and indirect effects of climate change on amphibian populations. - Diversity 2: 281–313.

Bókony, V. et al. 2017. Climate-driven shifts in adult sex ratios via sex reversals: The type of sex determination matters. - Philos. Trans. R. Soc. B Biol. Sci. 372: 20160325.

Brans, K. I. et al. 2018. Urban hot-tubs: Local urbanization has profound effects on average and extreme temperatures in ponds. - Landsc. Urban Plan. 176: 22–29.

Brooks, M. E. et al. 2017. glmmTMB balances speed and flexibility among packages for zero-inflated generalized linear mixed modeling. - R J. 9: 378–400.

Burraco, P. et al. 2020. Climate change and ageing in ectotherms. - Glob. Chang. Biol. 26: 5371–5381.

Campbell Grant, E. H. et al. 2016. Quantitative evidence for the effects of multiple drivers on continental-scale amphibian declines. - Sci. Rep. 6: 25625.

Carreira, B. M. et al. 2020. Heat waves trigger swift changes in the diet and life-history of a freshwater snail. - Hydrobiologia 847: 999–1011.

Ceballos, G. et al. 2015. Accelerated modern human-induced species losses: Entering the sixth mass extinction. 1: e1400253.

Chatfield, M. W. H. and Richards-Zawacki, C. L. 2011. Elevated temperature as a treatment for Batrachochytrium dendrobatidis infection in captive frogs. - Dis. Aquat. Organ. 94: 235–238.

Christensen, R. H. B. 2015. A tutorial on fitting Cumulative Link Mixed Models with clmm2 from the ordinal package.: 1–18.

Clarke, A. and Pörtner, H.-O. 2010. Temperature, metabolic power and the evolution of endothermy. - Biol. Rev. 85: 703–727.

Cohen, J. M. et al. 2017. The thermal mismatch hypothesis explains host susceptibility to an emerging infectious disease. - Ecol. Lett. 20: 184–193.

Cohen, J. M. et al. 2019. An interaction between climate change and infectious disease drove widespread amphibian declines. - Glob. Chang. Biol. 25: 927–937.

Courtney Jones, S. K. et al. 2015. Long-term changes in food availability mediate the effects of temperature on growth, development and survival in striped marsh frog larvae: Implications for captive breeding programmes. - Conserv. Physiol. 3: 1–12.

Crespi, E. J. and Warne, R. W. 2013. Environmental conditions experienced during the tadpole stage alter post-metamorphic glucocorticoid response to stress in an amphibian. - Integr. Comp. Biol. 53: 989–1001.

De Block, M. and Stoks, R. 2008. Compensatory growth and oxidative stress in a damselfly. - Proc. Biol. Sci. 275: 781–785.

Denver, R. J. 1997. Proximate mechanisms of phenotypic plasticity in amphibian metamorphosis. - Am. Zool. 37: 172–184.

Dournon, C. et al. 1984. Cytogenetic and genetic evidence of male sexual inversion by heat treatment in the newt Pleurodeles poireti. - Chromosoma 90: 261–264.

Fang, X. and Stefan, H. G. 2009. Simulations of climate effects on water temperature, dissolved oxygen, and ice and snow covers in lakes of the contiguous United States under past and future climate scenarios. - Limnol. Oceanogr. 54: 2359–2370.

Ferreira, V. and Chauvet, E. 2011. Synergistic effects of water temperature and dissolved nutrients on litter decomposition and associated fungi. - Glob. Chang. Biol. 17: 551–564.

Floyd, R. B. 1983. Ontogenetic change in the temperature tolerance of larval Bufo marinus (Anura: Bufonidae). - Comp. Biochem. Physiol. -- Part A Physiol. 75: 267–271.

Forrest, M. J. and Schlaepfer, M. A. 2011. Nothing a hot bath won’t cure: Infection rates of amphibian chytrid fungus correlate negatively with water temperature under natural field settings. - PLoS One in press.

Freitas, J. S. and Almeida, E. A. 2016. Antioxidant defense system of tadpoles (Eupemphix nattereri) exposed to changes in temperature and pH. - Zoolog. Sci. 33: 186–194.

Gardner, J. et al. 2016. Individual and demographic consequences of reduced body condition following repeated exposure to high temperatures. - Ecology 97: 786–795.

Geiger, C. C. et al. 2011. Elevated temperature clears chytrid fungus infections from tadpoles of the midwife toad, Alytes obstetricans. - Amphibia-Reptilia 32: 276–280.

Girish, S. and Saidapur, S. K. 2000. Interrelationship between food availability, fat body, and ovarian cycles in the frog, Rana tigrina, with a discussion on the role of fat body in anuran reproduction. - J. Exp. Zool. 286: 487–493.

Goldstein, J. A. et al. 2017. The effect of temperature on development and behaviour of relict leopard frog tadpoles. - Conserv. Physiol. 5: 1–8.

Gosner, K. L. 1960. A simplified table for staging anuran embryos larvae with notes on identification. - Herpetologica 16: 183–190.

Harkey, G. A. and Semlitsch, R. D. 1988. Effects of temperature on growth, development, and color polymorphism in the Ornate chorus frog Pseudacris ornata. - Copeia 4: 1001–1007.

Harvell, C. D. et al. 2002. Climate warming and disease risks for terrestrial and marine biota. - Science (80-.). 296: 2158–2162.

Hector, K. L. et al. 2012. Consequences of compensatory growth in an amphibian. - J. Zool. 286: 93–101.

Hettyey, A. et al. 2019. Mitigating disease impacts in amphibian populations: Capitalizing on the thermal optimum mismatch between a pathogen and its host. - Front. Ecol. Evol. 7: 1–13.

Hof, C. et al. 2011. Additive threats from pathogens, climate and land-use change for global amphibian diversity. - Nature 480: 516–519.

Holland, M. P. et al. 2007. Echinostome infection in green frogs (Rana clamitans) is stage and age dependent. - J. Zool. 271: 455–462.

IUCN 2021. International Union for Conservation of Nature. - https://www.iucnredlist.org/resources/summary-statistics in press.

Jonsson, B. and Jonsson, N. 2014. Early environment influences later performance in fishes. - J. Fish Biol. 85: 151–188.

Juráni, M. et al. 1973. Effect of stress and environmental temperature on adrenal function in Rana esculenta. - J. Endochrinology 57: 385–391.

Kilpatrick, A. M. et al. 2010. The ecology and impact of chytridiomycosis: an emerging disease of amphibians. - Trends Ecol. Evol. 25: 109–118.

Kitano, T. et al. 2012. Estrogen rescues masculinization of genetically female medaka by exposure to cortisol or high temperature. - Mol. Reprod. Dev. 79: 719–726.

Kozłowski, J. et al. 2004. Can optimal resource allocation models explain why ectotherms grow larger in cold? - Integr. Comp. Biol. 44: 480–493.

Krockenberger, A. et al. 2012. The limit to the distribution of a rainforest marsupial folivore is consistent with the thermal intolerance hypothesis. - Oecologia 168: 889–899.

Lambert, M. R. et al. 2018. Sexual and somatic development of wood frog tadpoles along a thermal gradient. - J. Exp. Zool. Part A Ecol. Integr. Physiol. 329: 72–79.

Lambert, M. R. et al. 2019. Molecular evidence for sex reversal in wild populations of green frogs (Rana clamitans). - Peer J. 7: e6449.

Laugen, A. T. et al. 2003. Latitudinal and temperature-dependent variation in embryonic development and growth in Rana temporaria. - Oecologia 135: 548–554.

Lindauer, A. L. et al. 2020. Daily fluctuating temperatures decrease growth and reproduction rate of a lethal amphibian fungal pathogen in culture. - BMC Ecol. 20: 1–9.

Lips, K. R. 2016. Overview of chytrid emergence and impacts on amphibians. - Philos. Trans. R. Soc. B Biol. Sci. 371: 20150465.

Lushchak, V. I. 2011. Environmentally induced oxidative stress in aquatic animals. - Aquat. Toxicol. 101: 13–30.

Marantelli, G. et al. 2004. Distribution of the amphibian chytrid Batrachochytrium dendrobatidis and keratin during tadpole development. - Pacific Conserv. Biol. 10: 173–179.

Martel, A. et al. 2013. Batrachochytrium salamandrivorans sp. nov. causes lethal chytridiomycosis in amphibians. - Proc. Natl. Acad. Sci. U. S. A. 110: 15325–15329.

McKechnie, A. and Wolf, B. 2019. The physiology of heat tolerance in small endotherms. - Physiology 34: 302–313.

McLeod, I. M. et al. 2013. Climate change and the performance of larval coral reef fishes: The interaction between temperature and food availability. - Conserv. Physiol. 1: 1–12.

McMahon, T. A. et al. 2014. Amphibians acquire resistance to live and dead fungus overcoming fungal immunosuppression. - Nature 511: 224–227.

Mikó, Z. et al. 2017. Age-dependent changes in sensitivity to a glyphosate-based pesticide in tadpoles of the common toad (Bufo bufo). - Aquat. Toxicol. 187: 48–54.

Mikó, Z. et al. 2021. Sex reversal and ontogeny under climate change and chemical pollution: are there interactions between the effects of elevated temperature and a xenoestrogen on early development in agile frogs? - Environ. Pollut. 285: 117464.

Monastersky, R. 2014. Biodiversity: Life a a status report. - Nature 516: 158–161.

Morand, A. et al. 1997. Phenotypic variation in metamorphosis in five anuran species along a gradient of stream influence. - Comptes Rendus l’Académie des Sci. - Ser. III - Sci. la Vie / Life Sci. 320: 645–652.

Murillo-Rincón, A. et al. 2017. Compensating for delayed hatching reduces offspring immune response and increases life-history costs. - Oikos 126: 565–571.

Narayan, E. J. and Hero, J.-M. 2014. Acute thermal stressor increases glucocorticoid response but minimizes testosterone and locomotor performance in the cane toad (Rhinella marina). - PLoS One 9: 1–6.

Nemesházi, E. et al. 2020. Novel genetic sex markers reveal high frequency of sex reversal in wild populations of the agile frog (Rana dalmatina) associated with anthropogenic land use. - Mol. Ecol. 29: 3607–3621.

Nemesházi, E. et al. 2021. Evolutionary and demographic consequences of temperature-induced masculinization under climate warming: the effects of mate choice. - BMC Ecol. Evol. 21: 16.

Niehaus, A. C. et al. 2006. Short- and long-term consequences of thermal variation in the larval environment of anurans. - J. Anim. Ecol. 75: 686–692.

O’Hanlon, S. J. et al. 2018. Recent Asian origin of chytrid fungi causing global amphibian declines. - Science (80-.). 360: 621–627.

Ogielska, M. and Kotusz, A. 2004. Pattern and rate of ovary differentiation with reference to somatic development in Anuran amphibians. - J. Morphol. 259: 41–54.

Ortiz-Santaliestra, M. E. et al. 2006. Influence of developmental stage on sensitivity to ammonium nitrate of aquatic stages of amphibians. - Environ. Toxicol. Chem. 25: 105–111.

Oswald, S. et al. 2008. Heat stress in a high-latitude seabird: effects of temperature and food supply on bathing and nest attendance of great skuas Catharacta skua. - J. Avian Biol. 39: 163–169.

Phuge, S. K. 2017. High temperatures influence sexual development differentially in male and female tadpoles of the Indian skipper frog, Euphlyctis cyanophlyctis. - J. Biosci. 42: 449–457.

Pierantoni, R. et al. 1983. Fat body and autumn recrudescence of the ovary in Rana esculenta. - Comp. Biochem. Physiol. -- Part A Physiol. 76: 31–35.

Pike, N. 2011. Using false discovery rates for multiple comparisons in ecology and evolution. - Methods Ecol. Evol. 2: 278–282.

Pounds, J. A. et al. 2006. Widespread amphibian extinctions from epidemic disease driven by global warming. - Nature 439: 161–167.

Retallick, R. W. R. and Miera, V. 2007. Strain differences in the amphibian chytrid Batrachochytrium dendrobatidis and non-permanent, sub-lethal effects of infection. - Dis. Aquat. Organ. 75: 201–207.

Richards-Zawacki, C. L. 2010. Thermoregulatory behaviour affects prevalence of chytrid fungal infection in a wild population of Panamanian golden frogs. - Proc. R. Soc. B Biol. Sci. 277: 519–528.

Ruiz-Garciá, A. et al. 2021. Sex differentiation in amphibians: Effect of temperature and its influence on sex reversal. - Sex. Dev. 15: 157–167.

Ruxton, G. D. and Beauchamp, G. 2008. Time for some a priori thinking about post hoc testing. - Behav. Ecol. 19: 690–693.

Scheele, B. C. et al. 2019. Amphibian fungal panzootic causes catastrophic and ongoing loss of biodiversity. - Science (80-.). 363: 1459–1463.

Scott, D. E. et al. 2007. Amphibian lipid levels at metamorphosis correlate to post-metamorphic terrestrial survival. - Oecologia 153: 521–532.

Spitzen-van der Sluijs, A. et al. 2016. Expanding distribution of lethal amphibian fungus Batrachochytrium salamandrivorans in Europe. - Emerg. Infect. Dis. 22: 1286–1288.

Squires, Z. E. et al. 2010. Compensatory growth in tadpoles after transient salinity stress. - Mar. Freshw. Res. 61: 219–222.

Stefan, H. G. et al. 2001. Simulated fish habitat changes in North American lakes in response to projected climate warming. - Trans. Am. Fish. Soc. 130: 459–477.

Stillman, J. H. 2019. Heat waves, the new normal: Summertime temperature extremes will impact animals, ecosystems, and human communities. - Physiology 34: 86–100.

Stoks, R. et al. 2006. Physiological costs of compensatory growth in a damselfly. - Ecology 87: 1566–74.

Stuart, S. N. et al. 2004. Status and trends of amphibian declines and extinctions worldwide. - Science 306: 1783–1786.

Sunday, J. M. et al. 2011. Global analysis of thermal tolerance and latitude in ectotherms. - Proc. R. Soc. B Biol. Sci. 278: 1823–1830.

Truebano, M. et al. 2018. Thermal strategies vary with life history stage. - J. Exp. Biol. 221: jeb171629.

Ultsch, G. R. et al. 1999. Physiology: Coping with the environment. - In: McDiarmid, R. W. and Altig, R. (eds), Tadpoles: the biology of anuran larvae. University of Chicago Press, Chicago, USA., pp. 189–214.

USEPA 2002. Methods for Measuring the Acute Toxicity of Effluents and Receiving Waters to Freshwater and Marine Organisms. - United States Environmental Protection Agency Office of Water (4303T).

Üveges, B. et al. 2016. Experimental evidence for beneficial effects of projected climate change on hibernating amphibians. - Sci. Rep. 6: 26754.

Van Rooij, P. et al. 2015. Amphibian chytridiomycosis: A review with focus on fungus-host interactions. - Vet. Res. 46: 1–22.

Voyles, J. et al. 2009. Pathogenesis of chytridiomycosis, a cause of catastrophic amphibian declines. - Science (80-.). 326: 582–585.

Voyles, J. et al. 2017. Diversity in growth patterns among strains of the lethal fungal pathogen Batrachochytrium dendrobatidis across extended thermal optima. - Oecologia 184: 363–373.

Wake, D. B. and Vredenburg, V. T. 2008. Are we in the midst of the sixth mass extinction? A view from the world of amphibians. - Proc. Natl. Acad. Sci. U. S. A. 105: 11466–73.

Wallace, H. and Wallace, B. M. N. 2000. Sex reversal of the newt Triturus cristatus reared at extreme temperatures. - Int. J. Dev. Biol. 44: 807–810.

Wedekind, C. 2017. Demographic and genetic consequences of disturbed sex determination. - Philos. Trans. R. Soc. B Biol. Sci. 372: 20160326.

Welbergen, J. A. et al. 2008. Climate change and the effects of temperature extremes on Australian flying-foxes. - Proc. R. Soc. B Biol. Sci. 275: 419–425.

Whiteley, S. L. et al. 2021. Temperature-induced sex reversal in reptiles: Prevalence, discovery, and evolutionary implications. - Sex. Dev. 15: 148–156.

Williams, C. M. et al. 2016. Biological impacts of thermal extremes: mechanisms and costs of functional responses matter. - Integr. Comp. Biol. 56: 73–84.

Woodhams, D. C. and Alford, R. A. 2005. Ecology of chytridiomycosis in rainforest stream frog assemblages of tropical Queensland. - Conserv. Biol. 19: 1449–1459.

Woodhams, D. C. et al. 2003. Emerging disease of amphibians cured by elevated body temperature. - Dis. Aquat. Organ. 55: 65–67.

Xu, Y. et al. 2021. Male heterogametic sex determination in Rana dybowskii based on sex-linked molecular markers. - Integr. Zool.: in press.

